# Targeted editing of *GTR1* and *GTR2* homologs directs the development of ideal low-seed and high-leaf glucosinolate oilseed mustard

**DOI:** 10.1101/2022.10.03.510517

**Authors:** Juhi Kumari, Avni Mann, Roshan Kumar, Akshay K. Pradhan, Deepak Pental, Naveen C. Bisht

**Author notes:** These authors contributed equally (Juhi Kumari, Avni Mann). Corresponding author: Naveen C. Bisht, Tele. **Author contributions:** N.C.B. conceived and planned the research work. J.K., A.M. and R.K. performed the experiments. N.C.B., D.P. and A.K.P. discussed the results and wrote the manuscript.

## Abstract

Glucosinolate content in the *Brassica* oilseed crops has been reduced to the globally accepted Canola quality levels, making the seed meal useful as a protein-rich animal feed. However, the overall lower glucosinolate content in such plants renders them vulnerable to pests. We report CRISPR/Cas9-based editing of glucosinolate transporter (*GTR*) family genes to develop ideal oilseed mustard lines with the desired low levels of seed glucosinolates while maintaining high glucosinolate levels in leaves and pod walls, without affecting the seed yield and seed quality traits. The low-seed and high-leaf glucosinolate mustard lines with improved nutritional and defense traits will raise its international trade potential both as oilseed and feed crop.

## Introduction

Glucosinolates (GSLs), ubiquitously present in the species belonging to the family *Brassicaceae*, provide defense against pests and pathogens^1,2,3^. However, the presence of high amounts of GSLs in the two major oilseed Brassica crops – *B. napus* (rapeseed) and *B. juncea* (mustard), and their hydrolysis products render the seed meal unpalatable to cattle and anti-nutritional to swine and poultry^4,5^. Over the last few decades, a priority breeding objective in the two oilseed *Brassica* crops has been to reduce the seed GSL content (SGC) to the Canola quality limit i.e., less than 30 μmoles g^−1^ seed dry weight (DW). However, such Canola quality rapeseed and mustard cultivars have low GSL content in all the plant organs, rendering the plants vulnerable to pests and pathogens^6,7^. The ideal lines should be – low GSL in the seeds, high GSL in the leaves and other plant parts.

GSLs are synthesized in the leaves and pod walls (source) and translocated to the seeds (sink) through the involvement of two related but distinct GSL transporters, GTR1 and GTR2^8^. In the model species *Arabidopsis thaliana*, null mutations in *GTR1* or *GTR2* provide some reduction in SGC, however, only the *gtr1gtr2* double mutant has SGC below the detection limits. TILLING mutagenesis^9^ and RNAi-suppression^10^ of the *GTR* gene families have been tested earlier in the allotetraploid *B. juncea* (AABB, 2n=36) for reducing the SGC. However, mustard lines containing SGC within the desired Canola quality levels have not been established till date. We report the development of an ideal GSL chemotype with low GSL in the seeds (sink) and high GSL in the source tissues (leaves and pod wall) by editing multiple *GTR1* and *GTR2* family homologs in oilseed mustard, without compromising the growth and development, and yield of the plants.

## Results and Discussion

Analysis of *GTR* gene families in our *B. juncea* cv. Varuna genome assembly^11^ showed the presence of six homologs of each of the *GTR1* and *GTR2* genes. To edit multiple *BjuGTR*s, we selected the most conserved region from the second exon and designed three guide RNAs (gRNAs) that can target a total of ten well-expressed *BjuGTR1* and *BjuGTR2* homologs (Fig. 1a). The three gRNA were cloned in a binary vector that contained the marker gene *bar* conferring resistance to herbicide ‘Basta’, and the *Cas9* gene to form the GTR1:GTR2(Ged) construct (Fig. 1b). The GTR(Ged) construct was used for *Agrobacterium*-mediated genetic transformation of *B. juncea* cv. Varuna which has a high SGC (134.44 ±5.01 μmoles g^−1^ DW). A total of 37 independent transgenic events (T0) were generated. T1 seeds of each event were analyzed for their SGC content, which ranged from 6.21 to 145.88 μmoles g^−1^ DW (Fig. 1c). A significant reduction of SGC at par with the Canola quality standard was observed in T1 seeds of the 30 events (Dataset S1).

**Figure 1.**
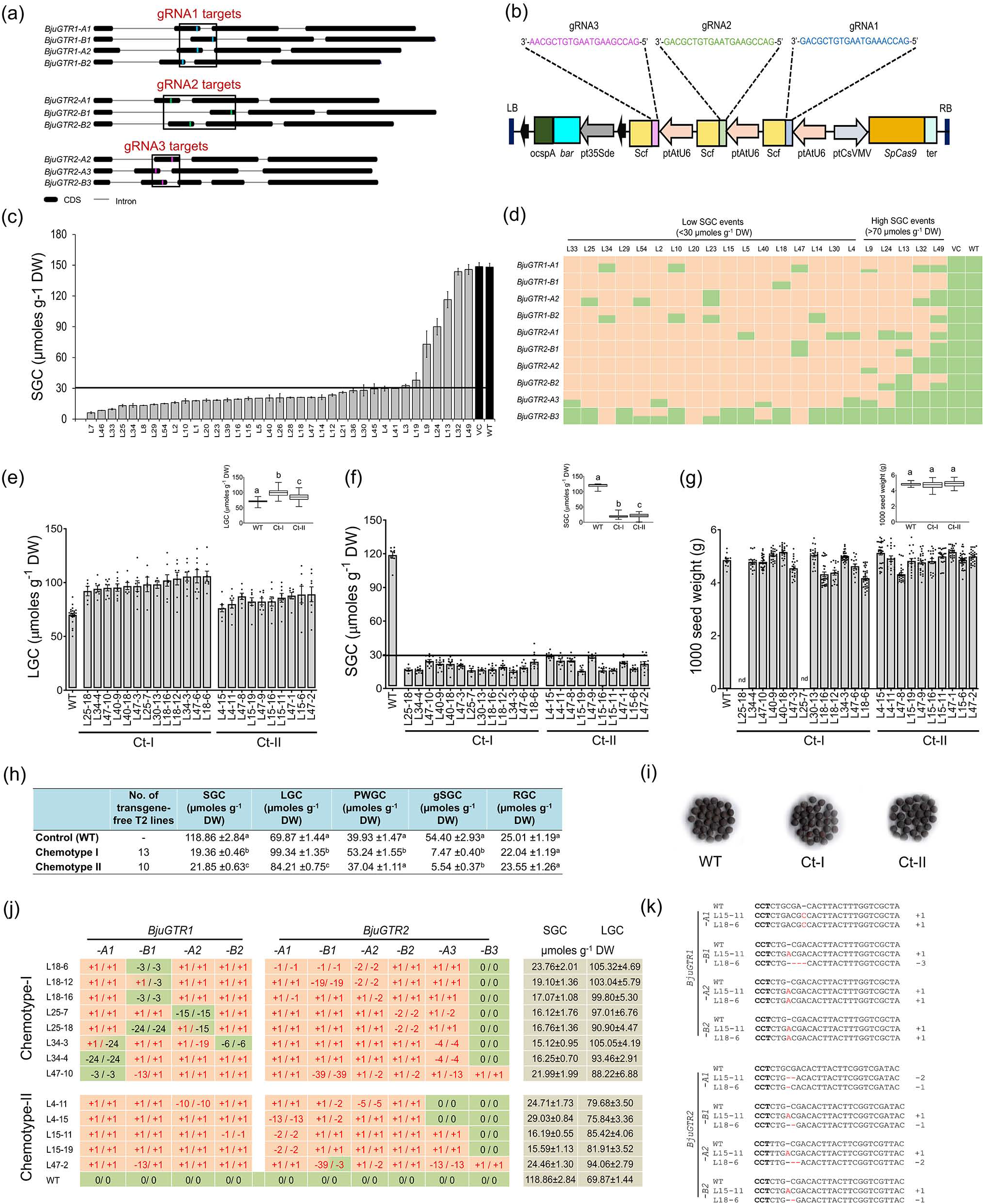
Editing of *GTR1* and *GTR2* homologs and development of transgene-free mustard lines containing low-seed, high-leaf glucosinolate content. **(a)** Genomic locations of gRNA target sites on the ten *BjuGTR* homologs. **(b)** T-DNA map of GTR1:GTR2(Ged) construct containing gRNA1 (blue), gRNA2 (green) and gRNA3 (violet). **(c)** Mean SGC in T1 seeds of 37 independent T0 events, vector control (VC), and wild-type (Varuna) plants; the Canola quality limit (30 µmoles g^−1^ DW) is marked with a line. **(d)** Analysis of the CRISPR/Cas9-induced mutations in ten *BjuGTR* homologs of T0 events with low and high SGC; editing in both the alleles of *BjuGTR* homolog have been marked in orange, a single edited allele in orange/green, and both alleles unedited in green color. **(e)** Leaf GSL content (LGC) at the post-bolting stage, and **(f)** mature seed GSL content (SGC) of 23 transgene-free *BjuGTR*-edited lines in the T2 generation. **(h)** Summary of the mean GSL content in mature seeds (SGC), leaves (LGC), pod walls (PWGC), developing green seeds (gSGC), and roots (RGC) of the lines belonging to chemotypes Ct-I (13 lines) and Ct-II (10 lines). **(g)** Seed size (measured as 1000 seed weight), and **(i)** seed shape of the representative transgene-free *BjuGTR*-edited lines. **(j)** Analysis of the genetic zygosity of *BjuGTR* mutations in 14 representative low SGC lines belonging to the two chemotypes; the length of the insertion (+) or deletion (-) observed in the two alleles of each *BjuGTR* gene with respect to the wild-type alleles (0 / 0) is provided. **(k)** Genotype of mutations in well-expressed *BjuGTR1* and *BjuGTR2* homologs at the target sites belonging to a representative Ct-I and Ct-II plant.

We studied the spectrum of CRISPR/Cas9-induced mutations in the ten *BjuGTR* target sites of 17 events showing SGC <30 μmoles g^−1^ DW in T1 seeds (Fig. 1d). The use of three gRNAs in a single construct provided >84% mutation efficiency in all the target *BjuGTR* homologs, except *BjuGTR2-B3* (Dataset S2). More than 90% of the mutations were single nucleotide insertions or deletions (indels) leading to frameshift null mutations in *BjuGTR1* and *BjuGTR2* homologs (Dataset S3). We also analyzed the mutation profile in *BjuGTR* homologs of five events containing high SGC (>70 μmoles g^−1^ DW) and compared this with the low SGC events (Fig. 1d). In the high SGC events, a few *BjuGTR2* homologs either remained unedited or contained short deletions in the multiples of 3 bp (Dataset S3), which might have maintained the functionality of these genes.

We self-pollinated the low SGC events (T0) and identified the marker and *Cas9*-free plants in the T1 generation by their sensitivity to ‘Basta’ and PCR analysis (Dataset S4). A total of 23 independent transgene-free lines, having low SGC in the T2 seeds were selected (Table S5) and self-pollinated to obtain the T2 generation plants.

We further checked GSL accumulation in the leaf (post-bolting stage) and the pod (silique) walls in 8-10 T2 generation plants of each of the 23 low SGC lines. The leaf GSL content (LGC) in the low SGC lines was found to be ranging from 75.84 to 105.32 μmoles g^−1^ DW which was higher than that observed in the wild-type plants (69.87 ±1.44 μmoles g^−1^ DW) (Fig. 1e,h; Dataset S6). Thirteen out of 23 tested lines accumulated significantly higher LGC (99.34 ±1.35 μmoles g^−1^ DW) compared to the wild-type plants and were categorized as chemotype-I (Ct-I). The remaining 10 lines showed marginally higher LGC (84.21 ±0.75 μmoles g^−1^ DW) compared to the control plants and were categorized as Ct-II. In addition, the pod wall GSL content (PWGC) of the Ct-I lines (53.24 ±1.55 μmoles g^−1^ DW) was also higher than that in the Ct-II lines (37.04 ±1.11 μmoles g^−1^ DW) and the wild-type plants (39.93 ±1.47 μmoles g^−1^ DW) (Fig. 1h; Dataset S7). The developing green seeds of low SGC events were also tested for their GSL content (gSGC), which was found to be significantly lower than in the wild-type seeds (Fig. 1h), clearly showing impaired GSL transport from the pod walls to the developing seeds. The root GSL content (RGC) of the two GSL chemotypes was comparable to that of the wild-type plants (Fig. 1h). The T3 generation seeds of all the selected lines accumulated mean SGC ranging from 15.12 to 29.03 μmoles g^−1^ DW which is below the Canola quality threshold (Fig. 1f,h, Table S8).

The *BjuGTR*s mutation spectrum in 23 transgene-free low SGC-high LGC lines was analyzed in the T2 generation plants. The mutations in the ten *BjuGTR* homologs were mostly in true biallelic homozygous conditions (Fig. 1j,k; Dataset S9). All the 23 low SGC lines contained loss-of-function mutations in a minimum of four or more homologs of *BjuGTR2* (*2-A1, 2-B1, 2-A2* and *2-B2*). However, an interesting difference was observed between the lines belonging to the Ct-I and Ct-II categories (Fig. 1j,k). While the lines belonging to chemotype Ct-II had loss-of-function mutations in all the four well-expressed *BjuGTR1* homologs (*1-A1, 1-B1, 1-A2*, and *1-B2*), the lines belonging to the category Ct-I, with significantly higher over-accumulation of LGC and PWGC, contained at least one or two functional *BjuGTR1* homolog/s. The functional *BjuGTR1* homologs could be helping in maintaining an optimal GSL distribution in source organs in the Ct-I plants.

We also studied the 23 low SGC, high LGC mustard lines grown in soil in a containment net-house during the mustard growing season 2021-22 for their growth and development and yield-related traits that included seed germination, plant height, pollen viability, seed set, and 1000 seed weight, and seed quality parameters that included oil content, protein content, and fatty acid composition. The values for all these traits in the low SGC, high LGC lines were found to be comparable to that of the wild-type plants (Figure 1g,i; Dataset S10).

In *Arabidopsis*, the loss-of-function of both *GTR1* and *GTR2* genes resulted in a reduction in the seed weight by 12±3% and total seed yield by 26±8% compared to the wild-type plants^8^. In the related allopolyploid *B. napus* (AACC), editing of only the *BnaGTR2* homologs reduced the accumulation of GSLs in the sink (seeds) as well as in the source organs (leaf and siliques)^12,13^. In addition, the seed size, amino acid accumulation, sugar content and fatty acid composition were also adversely affected^12^. The strategy of simultaneous editing of both *GTR1* and *GTR2* homologs may also work in rapeseed where editing of only *GTR2* homologs did not provide the ideal chemotype.

The conventionally bred Canola quality lines of rapeseed have become increasingly susceptible to generalist pests in Europe and elsewhere^6,7^ leading to increased use of pesticides for crop protection. We earlier developed mustard lines with low SGC using both conventional breeding^14^ and RNAi-suppression of *BjuMYB28* genes^15^. However, these low SGC lines, due to reduced GSL accumulation throughout the plant, were found to be vulnerable to heavy damage by termites, generalist pests, birds, and herbivores. The transgene-free *BjuGTR*-edited mustard lines developed in this study contain low SGC while providing a concomitant over-accumulation of GSL in the leaves and pod walls, which may even improve the plant defense. Our approach will contribute immensely towards developing Canola quality rapeseed-mustard cultivars thus raising their oil- and seed-meal quality in the global market as well as dependency on the agrochemicals.

## Acknowledgments

The work was supported by the DBT (India) grant BT/HRD-NBA-NWB/38/2019-20(10) to N.C.B. Financial support from UGC (India), CSIR (India), and DBT (India) to J.K., A.M., and R.K. respectively, is acknowledged. The central facilities of NIPGR are also acknowledged.

## Supplementary Information

The Supplementary Information includes a Materials and Methods section and a supporting Dataset containing eleven Tables (as excel dataset).

**Dataset Table S1:** Estimation of fractions and total SGC (µmoles g^−1^ DW) in the T1 seeds of *BjuGTR*-edited events and the wild-type (WT) Varuna and vector control (VC) plants.

**Dataset Table S2:** Mutation frequency and average mutation efficiency recorded in different *BjuGTR1 and BjuGTR2* homologs in 22 independent *BjuGTR*-edited T0 events.

**Dataset Table S3:** Analysis of CRISPR/Cas9-induced mutation in the *BjuGTR* homologs in 22 selected *BjuGTR*-edited lines in the T0 generation.

**Dataset Table S4:** Summary of *Cas9*-free (transgene-free) T1 plants obtained for each transformation event derived from GTR1:GTR2(Ged) construct.

**Dataset Table S5:** Estimation of various GSL fractions and the total GSL content (µmoles g^−1^ DW) in T2 seeds of *BjuGTR*-edited lines along with the wild-type (WT) Varuna and vector control plants.

**Dataset Table S6:** Estimation of various GSL fractions and the total GSL content (µmoles g^−1^ DW) in the young leaf of 23 transgene-free low SGC lines and the wild-type (WT) Varuna plants.

**Dataset Table S7:** Estimation of various GSL fractions and the total GSL content (µmoles g^−1^ DW) in the pod walls (PWGC) of 23 transgene-free low SGC lines and the wild-type (WT) Varuna plants.

**Dataset Table S8:** Estimation of various GSL fractions and the total GSL content (µmoles g^−1^ DW) in mature T3 seeds (SGC) of 23 transgene-free low SGC lines and wild-type (WT) Varuna plants.

**Dataset Table S9:** Analysis of CRISPR/Cas9-induced mutations in the *BjuGTR* homologs in the T2 generation of the selected transgene-free *BjuGTR*-edited lines.

**Dataset Table S10:** Seed quality and yield-related parameters (mean ± standard error) of mature T3 seeds of 23 low SGC lines and wild-type (WT) Varuna seeds estimated using near-infrared spectroscopy (NIRS).

**Dataset Table S11:** List of primers used in this study.

## Materials and Methods (Online)

### Designing gRNAs and generation of CRISPR/Cas9-based editing construct/s

Full-length genomic sequences of *GTR1* (*BjuGTR1*) and *GTR2* (*BjuGTR2*) homologs from the genome assembly of *B. juncea* cultivar Varuna were used as queries to identify the 20 nt gRNAs using CRISPR-P v2.0 (http://crispr.hzau.edu.cn/CRISPR2/). Three gRNAs targeting all the expressed *BjuGTR* homologs were selected based on maximum on-score and minimum off-targets while containing 5’ G and 3’ NGG as PAM sequences for the *Streptococcus pyogenes* Cas9. To generate a sgRNA scaffold, the 20 nt seed sequence of gRNA was introduced between the AtU6-26 promoter and the scaffold in a two-step PCR reaction strategy, using customized PCR primers (Table S11). In the first step, two PCR reactions were performed to amplify the ‘promoter-gRNA’ and ‘gRNA-scaffold’ fragments, independently. In the second step, the two fragments were linked through another round of overlapping PCR to generate the AtU6-26 promoter:gRNA:scaffold expression cassette (Fig. S2).

The *S. pyogenes* wild-type *Cas9* gene (*SpCas9*), driven by the constitutively expressing Cassava Mosaic Virus (CsVMV) promoter was cloned into the binary vector pZP200debar^15^ containing *bar* gene conferring resistance to herbicide Basta (Bayer, Germany) as a selectable marker (within the ‘lox’ tandem repeats) to develop the pZP200debar::SpCas9 binary vector. All the three sgRNA fragments were cloned at the appropriate restriction sites in the pZP200debar::SpCas9 vector to develop the GTR1:GTR2(Ged) construct (Fig. 1b).

### Genetic transformation of *B. juncea* and development of *GTR-*edited lines

Genetic transformation of a high glucosinolate *B. juncea* cv. Varuna was performed as per the protocol established in the laboratory^15^. *B. juncea* seeds were cleaned by washing with a few drops of detergent (Teepol), followed by treatment with 70% ethanol for 2 min, and surface sterilized with 0.05% HgCl_2_ for 10 min. Seeds were thoroughly washed with sterile water at least five times after every treatment. Sterilized seeds were then inoculated on the basal MS media and kept in a culture room under a 10 hrs light/14 hrs dark cycle and a constant temperature of 23 ±1 °C. Hypocotyls of 5 days old seedlings were cut into ∼1 cm pieces and used as explants, which were precultured for 24 hrs at 110 rpm shaking in liquid MS medium containing 1 mg l^−1^ each of NAA (1-Naphthalene Acetic Acid) and BAP (6-Benzylaminopurine) referred to as N1B1. *Agrobacterium tumefaciens* strain GV3101 carrying the desired construct was grown on a YEB liquid medium containing appropriate antibiotics (10 mg l^−1^ Rifampicin, 10 mg l^−1^ Gentamycin and 50 mg l^−1^ Spectinomycin). Bacterial cells were harvested at OD_600_ of 0.5-0.6 and re-suspended in liquid N1B1 medium; the bacterial suspension was adjusted to the final OD_600_ to 0.3. Precultured explants were then incubated with the bacterial suspension in dark for 30 min and co-cultivated in liquid N1B1 medium at 23 ±1 °C, 110 rpm for 16-24 hrs. After co-cultivation, explants were washed with liquid N1B1 medium supplemented with 200 mg l^−1^ Augmentin to remove excess *Agrobacterium* cells. Explants were plated on MS agar medium supplemented with 1 mg l^−1^ each of NAA and BAP, 200 mg l^−1^ Augmentin, and 10 mg l^−1^ herbicide Basta (active ingredient phosphinothricin; Bayer AG, Germany). Regenerated shoots were transferred to the rooting media (2IBA) i.e., MS agar medium containing 2 mg l^−1^ IBA (Indole-3-acetic acid) along with 200 mg l^−1^ Augmentin and selection agent, Basta. The well-rooted T0 transformed plants were planted directly into the soil in a containment net-house under field conditions at NIPGR during the mustard growing season (Nov-Mar), as per the biosafety guidelines prescribed by the Department of Biotechnology, Government of India. T0 transformants were confirmed using Basta spray (200 mg l^−1^ phosphinothricin). The unopened flower buds of T0 plants were bagged to harvest the self-pollinated T1 seeds at maturity, which were used to identify the Basta resistant (transgenic) and Basta sensitive (*Cas9*-free or transgene-free) plants in the T1 generation. The Basta-sensitive T1 plants were self-pollinated to produce the T2 seeds.

### Isolation of genomic DNA and mutation screening

Genomic DNA was isolated from the young leaves for mutation analysis using the CTAB method. Approx. 200 bp of the genomic sequence flanking the target site of each *BjuGTR* homolog was amplified using the homolog-specific primers (Table S11) and ExTaq DNA polymerase (TaKaRa Bio Inc.). PCR product was gel-eluted and subjected to Sanger sequencing. The chromatogram file of each edited line was compared with that of the wild-type plant using the ICE analysis tool (Synthego Corporation, USA) to identify the mutations in the target genes. The edited and wild-type sequences were aligned using different modules of the DNASTAR software (Lasergene, USA).

The percentage of plants showing editing of the target gene was used to calculate the editing frequency. Mutation efficiency for each gene was calculated based on the percentage of the cell population having insertions or deletions, as analyzed by the software. The allelic status of the target gene in the edited lines was categorized into biallelic homozygous (BHo), biallelic heterozygous (BHt), monoallelic (Mo) and chimeric (Chi).

### Identification of transgene-free *BjuGTR-*edited lines

Around 40-50 T1 seeds from each independent transformation event were grown in a containment net-house and tested for their sensitivity/resistance to herbicide Basta (200 mg l^−1^; active ingredient phosphinothricin) by painting a few young leaves of 3-4 week-old plants with the herbicide. Segregation of T1 progeny for Basta resistant (Cas9-containing) and sensitive (Cas9-free) phenotypes were recorded and the copy number of the inserted T-DNA was determined using the *Chi*-square analysis. Basta-sensitive transgene-free lines were confirmed for the absence of the *Cas9* and selectable marker (*bar*) genes by PCR analysis (Table S12; Fig. S4). Basta-sensitive (or Cas9-free) T1 plants were multiplied by self-pollination to obtain homozygous mutations for the *BjuGTR* homologs in the T2 generation.

### Glucosinolate estimations

Total seed GSL content and various seed quality parameters in mature seeds were estimated by near-infrared reflectance spectroscopy (NIRS-DS 2500, FOSS, Denmark). The total and component GSLs from different tissue types, namely seeds, leaves, green pods, and developing seeds were determined using high-pressure liquid chromatography, as described earlier^15^. Briefly, glucosinolates were extracted using 10-20 mg of freeze-dried sample in 70% methanol, containing 50 μM sinalbin as the internal standard. Desulfation of glucosinolates was performed overnight using purified sulfatase (25 mg ml^−1^; Sigma-Aldrich) on a DEAE Sephadex-A25 column. Desulfo-glucosinolates were eluted with 1 ml of HPLC-grade water and 30 μl of eluent was analyzed using a Shimadzu CLASS-VP V6.14 HPLC machine and Shim-pack GIST C18 reverse-phased column (4.6 × 250 mm; 5 μm i.d.) through a gradient of water (solvent A) and 1% to 19% (v/v) acetonitrile (solvent B) for 25 min with a flow rate of 1 ml min^−1^ with an oven temperature of 35 °C and detection was made at 229 nm.

Glucosinolate concentration (µmoles g^−1^ DW) was determined by identifying the substrate peak of known glucosinolates and referencing it with the known internal standard peaks. For internal standard sinalbin, the response factors (RF) used for quantifying aliphatic, benzenic, and indolic glucosinolates were 2, 0.5, and 0.5 respectively^15^. The total GSL content and profiles in T0 and T1 plants were estimated in 3-4 replications. For each low SGC event, 8-10 independent T2 progeny (transgene-free) were analyzed for determining the mean glucosinolate concentrations and profiles in mature seeds, flag leaf (at 50% flowering stage), developing green seeds, and pod walls (14-21 days post anthesis) and roots (one-month-old). Data were checked for normal distribution and homogeneity of variance through Shapiro-Wilk and Levene’s test respectively in SPSS. One-way ANOVA was performed using the Tukey’s/Games-Howell post hoc test in SPSS and P-value <0.05 was considered significant. Box plots were plotted using GraphPad Prism 9.

## Notes

**Conflict of interest:** The authors declare that no conflict of interest exists.

### Competing Interest Statement

The authors have declared no competing interest.

